# No evidence for extrafloral nectaries in *Erythranthe angulosa*

**DOI:** 10.64898/2026.08.28.747905

**Authors:** Sylvie Martin-Eberhardt, Paige Smith, Madison Plunkert

## Abstract

Extrafloral nectaries (EFNs) are a widespread plant defense mutualism trait and are highly convergent, appearing in hundreds of plant lineages worldwide. Here we investigate a report of possible EFNs in *Erythranthe angulosa*, a recently-described California wildflower. We integrate field observations, insect bioassays, an induction experiment, and microscopy to test for signatures of EFN function, finding no evidence that the distinctive axillary swellings produced by *E. angulosa* function as EFNs. We also uncovered two distinct morphs at the type locality of *E. angulosa* that diverge in the number of axillary swellings produced, as well as other shoot architecture traits such as stem thickness, leaf size, and branch number. Although the axillary swellings appear to not function as EFNs, they remain a compelling morphological variant within the yellow monkeyflowers that may perform storage or another unknown function.

---

Extrafloral nectaries (EFNs) are one of many strategies plants employ to avoid herbivory. This highly convergent, widespread defense mutualism is characterized by plants secreting nectar on their leaves and stems that attracts predatory insects including ants, wasps, or spiders whose presence ultimately reduces herbivory (Bentley 1977). EFNs have evolved in at least 457 angiosperm and pteridophyte lineages (Weber and Keeler 2013) and have arisen in diverse plant tissues (Marazzi et al. 2013a). Our knowledge of EFN diversity is continuously growing, with new reports of EFN-bearing species each year, bringing us closer to a complete picture of the phylogenetic breadth of this trait. While genetic model systems such as *Populus trichocarpa* and *Gossypium hirsutum* produce EFNs, we still lack an easily transformable EFN-bearing system, preventing large-scale genetic work on this widespread, convergent, and paradigmatic mutualistic trait.

EFNs are hugely variable in their anatomy and physiology, making investigations of diverse EFN-bearing lineages key to achieving a complete picture of this ecologically important trait. Across lineages, EFNs have been reported on leaves, petioles, stipules, stems, pedicels, and outer floral organs such as bracts (Marazzi et al. 2013a). Along with occurring in a wide range of tissue types, EFNs are also hugely variable in their morphology, ranging from highly ‘individualized’ secretory tissue to cryptic, nondifferentiated regions, as well as glandular trichomes (Marazzi et al. 2013b). In addition to distinct, discrete glands, EFNs also occur as many small, discontinuous nectar-secreting regions scattered over a large area and can range from superficial to deep within surrounding tissue (Nepi 2007). They also vary in their vascularization and phenology, with many EFNs being most active on young tissue (Nepi 2007; Calixto et al. 2021). Thus, investigation of EFNs across phylogenetically diverse systems can reveal the developmental, anatomical, and physiological mechanisms by which this defense mutualism trait has repeatedly evolved across flowering plants and ferns.

One newly described monkeyflower species, *Erythranthe angulosa*, has been reported to produce axillary swellings (Nesom and Berger 2020). Nesom and Berger suggest that these swellings perhaps function as EFNs (2020). *E. angulosa* is a member of the yellow monkeyflower species complex (*Erythranthe* sect. *Simiola*, (Barker et al. 2012), but see Lowry et al. 2019), a genetically diverse section showing local adaptation to herbivory (Popovic and Lowry 2020) and other environmental variables (Hall and Willis 2006; Lowry and Willis 2010; DeMarche et al. 2016). They produce a diverse array of structures at the leaf axils, including flowers, stolons, flowering branches, bulbils, and rhizomes (Moody et al. 1999; Baker and Diggle 2011; Friedman et al. 2015; Coughlan et al. 2021). With emerging tools for genetic and genomic analysis (Lovell et al. 2025; Stanley and Lowry 2025) and a long history of evolutionary ecology studies (Clausen and Hiesey 1958; Vickery 1978), monkeyflowers offer a powerful system for understanding how morphological diversity evolves and influences ecological interactions.

Verified production of EFNs in the genetically tractable yellow monkeyflower complex would offer opportunities to study EFN development, physiology, biochemistry, and ecology in this model system for evolutionary ecology and genetics. Additionally, an EFN-producing monkeyflower would be the first report of EFNs in Phrymaceae, further expanding the degree of convergence known in this trait across the plant tree of life. In this study, we investigate these axillary structures using field observations, laboratory experiments, and microscopy to test whether they function as EFNs.

## Methods

### Field Observations

We surveyed the plants growing in and around the type locality (36.467°N, 118.134°W) as reported in Nesom & Berger (2020) from June 30th to July 2nd, 2024. We checked plants for axillary swellings and recorded variation in the morphology and size of the swellings. To test for extrafloral nectar secretion, we bagged individual stems overnight, excluding EFN visitors who might remove nectar and reducing evaporation to maximize the chances of detecting secreted nectar droplets (Díaz-Castelazo et al. 2005). The following morning, we used glucose test strips (Bartovation, Queens, New York, USA) to test for sugar content in secreted fluid.

We also checked all plants for herbivory and categorized herbivory according to type (e.g. insect chewing herbivore, larger mammalian herbivore, etc), and collected potential EFN visitors and arthropod herbivores as voucher specimens. We observed no potential pollinators. In addition to field observations of ant or other arthropod visitation to EFNs, we buried several greenhouse-grown progeny (accession 1, see Germplasm Sampling below for accession details) in pots near the base of peony plants at Kellogg Biological Station (42.4056, -85.4024) such that the pot was flush with the mulch substrate. The peony bushes were in bud and were heavily visited by ants. Ants can be generalist visitors of a diversity of EFN-bearing plants (Aranda-Rickert et al. 2014), making any visitation or lack thereof informative. We observed several times throughout the day if ants were visiting the monkeyflower plants and if so, what region of the plant.

### Germplasm Sampling

We harvested a single ripe fruit from an individual that we identified in the field as *E. angulosa* due to its abundant axillary swellings for additional laboratory experiments, which we selfed for two generations in the lab and labelled ‘accession 1.’ No other clearly identifiable *E. angulosa* individuals bore mature seed at the time of collection. We also collected ∼100 mL of soil from beneath these plants for subsequent germination in the greenhouse. From this seed bank soil, we germinated 77 yellow monkeyflower individuals that included two distinct morphs: 51 large, highly branched plants and 26 short, large-leaved plants with simpler architecture (e.g. Fig. 3). We named these the “robust morph” and “thin-stemmed morph” respectively. To capture size variation within and between morphs, we selected the two tallest and two shortest plants of each morph grown from the seed bank for the subsequent induction experiment.

### Test for Genetic and Environmental Variation

Because EFNs from other systems have been demonstrated to upregulate nectar production in response to herbivory, both real and simulated (Koptur 1990; Heil et al. 2001; Kwok and Laird 2012; Hernandez-Cumplido et al. 2016) and even increase the number and size of EFN structures (Mondor and Addicott 2003; Pulice and Packer 2008; Yamawo and Suzuki 2018), we conducted an induction experiment with three treatments shown to increase nectar production or defense investment in other systems: Jasmonic Acid (JA), simulated herbivory (clipping) and silica-containing soil amendments. JA and simulated herbivory are well-demonstrated to increase investment in EFNs (Koptur 1990; Heil et al. 2001; Mondor and Addicott 2003; Pulice and Packer 2008; Kwok and Laird 2012; Hernandez-Cumplido et al. 2016; Yamawo and Suzuki 2018), and silica can interact with damage to increase EFN investment (Gowton et al. 2025) and is known to induce other herbivore defenses (Acevedo et al. 2021). We crossed these treatments with the families described above (accession 1, four families of the robust morph, and four families of the thin-stemmed morph). Plants were germinated after 14 days of stratification at 4°C and then grown in a chamber (Conviron Gen 2000, Winnipeg, Manitoba, Canada) on 14 light / 10 hour dark cycle with 28.3°C days (10% humidity) and 15.5°C nights (35% humidity), replicating July conditions at the type locality. All plants were fertilized weekly with half strength Hoagland’s solution (Hoagland’s No. 2 Basal Salt Mixture, Plant Cell Labs, Washington, DC, USA).

Treatments were imposed approximately every two weeks starting six weeks after seeds were removed from cold stratification. In addition to the control treatment, the clipping treatment had one third of all leaves removed with scissors and the JA treatment was sprayed (separately from other plants) with 1 mM jasmonic acid dissolved in 0.25% ethanol. This concentration of jasmonic acid has previously been shown to induce morphological response without detrimental effects in closely related *Erythranthe guttata* (Toll et al. 2026). The soil amendment treatment was grown in 9 parts soilless peat-based potting media (SureMix^TM^, Michigan Grower Products, Galesburg, Michigan, USA), 6 parts crushed granite (Yard Elements, Fallbrook, California, USA), and 1 part large grit silica sand (Asluru, Nanjing, Jiangsu, China). All other plants were grown in 100% SureMix^TM^. Three months after the treatments were assigned and applied, we visually inspected plants for signs of nectar production including testing for the presence of glucose using test strips, quantified the number of axillary swellings per plant, counted the number of branches, and measured the length of the longest leaf (excluding the petiole) and the stem thickness (at the midpoint of the lowest node accessible by a caliper).

All statistical analyses were performed in R Studio version 4.5.2 using the *glmmTMB* package (Brooks et al. 2017). All models of axillary swelling number were fit with a negative binomial distribution due to overdispersion. To quantify variation in axillary swelling production attributable to genetic (family) and environment (treatment) factors, we created an intercept-only model with family and treatment as random intercepts, and used the ‘icc’ function in the *performance* package (Lüdecke et al. 2021). To test for differences in axillary swellings among the two morphs and accession 1, we fit a negative binomial model with phenotype as a fixed effect. To test for morph differences in longest leaf length and stem thickness, we logged and scaled these variables and fit gaussian models with morph as a fixed effect and family as a random effect. For the morph difference in number of stems, we fit a negative binomial model due to overdispersion.

### Laboratory Observations of Crickets

As tissues rich in sugar and starch (Paul and Mitra 2024), EFNs can be targeted by herbivores (Gish et al. 2015). To test whether the axillary swellings are selectively herbivorized by a generalist herbivore, we starved 19 tropical house crickets (*Gryllodes sigillatus*) overnight before placing them individually with one intact branch from accession 1 plants in 150-mm petri dishes that were humidified with damp filter paper. After four days, we noted which parts of the branch had been damaged by cricket herbivory.

### Microscopy

All plants grown for microscopy analyses were second-generation inbred lines of accession 1 and were grown in growth chambers (model FXC-19; Bio Chambers Inc., Winnipeg, Manitoba, Canada) in a 16-hour photoperiod with a light intensity of 460 µmol m^-2^ s^-1^ photosynthetic active radiation, 22°C day and 18°C night temperatures, and 60% relative humidity.

We selected two stems with axillary swelling for analysis with scanning electron microscopy (SEM). We mounted fresh tissue on PELCO tabs affixed to aluminum stubs and secured with carbon tape. We imaged them immediately with a Hitachi TM3030 tabletop scanning electron microscope (Hitachi High-Tech Corporation, Tokyo, Japan). Because some EFNs secrete nectar through stomata (Nepi 2007) and floral nectar in monkeyflower is secreted through stomata (Zhai et al. 2025), we checked for a higher density of stomata on the axillary swellings.

For light microscopy, we cut transverse cross-sections of 3 axillary swellings and a longitudinal section of 1 axillary swelling using a double-edged razor blade. We stained fresh sections for several seconds in 0.02% Toluidine Blue O solution, rinsed in deionized water, and wet-mounted sections in water. We immediately viewed sections with an AmScope compound light microscope (AmScope, Irvine, California, USA) and imaged them with a Nikon microscope camera and Nikon BR software (Nikon, Tokyo, Japan).

## Results

### Field Observations

We located 3 plants with the distinct axillary swellings diagnostic for *E. angulosa* (Fig. 1A) growing from a small seep consisting of a single crevice in a granite wall. Additional plants were growing in an adjacent small indentation in the rock and did not appear to have axillary swellings. We also observed many seedling plants growing beneath accession 1 and neighboring plants; these seedlings appeared too young to observe the presence or absence of an EFN.

**FIG. 1.**
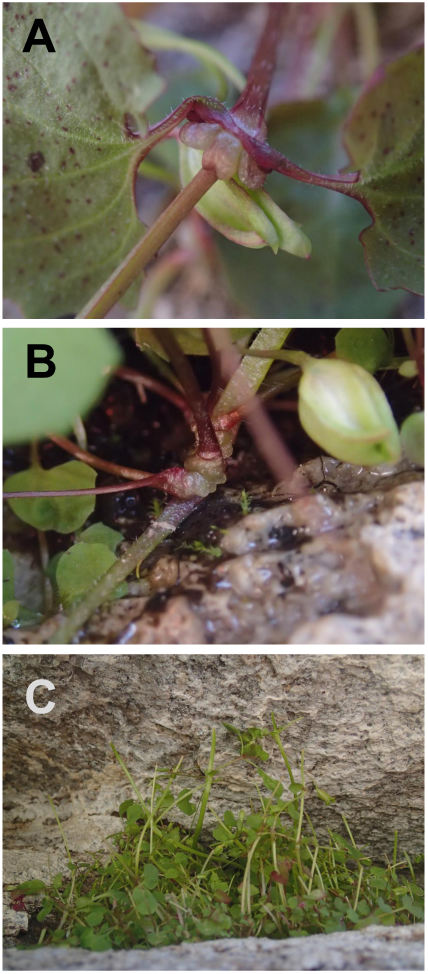
Observations of Erythranthe angulosa in the field including (A) a typical axillary swelling (B) an axillary swelling with extra lobing demonstrating the range of variation of this feature and (C) Extensive herbivory characteristic of mammals (entire stem consumed) in adjacent monkeyflower without developed axillary swellings. Photo credits: SME

Axillary swellings on these plants took a range of forms, with variation in the number of lobes, whether lobes met in a visible cleft, and whether swelling appeared to include branches, petioles, or pedicels. Swellings usually occurred on nodes bearing flowers or fruits and showed expansion of the base of the pedicel. We observed in both the field as well as subsequent greenhouse-grown individuals that axillary swellings were particularly conspicuous on branches that hung down below the root-shoot interface of the plant, e.g. down the side of the cliff or over the edge of a pot. After overnight bagging, we detected no glucose production in the field using glucose test strips, nor fluid production specific to the axillary swellings.

In the field, we observed extensive mammalian herbivory on the small plants growing in the adjacent indentation, characterized by clipping of the entire main stem of the plant (Fig. 1C). In contrast, accession 1 and surrounding plants growing in the crevice had only slight mammalian herbivory, although this may be attributable to herbivores’ access to the steep granite cliff rather than plant defense by insect bodyguards recruited by EFNs. We observed low insect herbivory overall, and recorded only a small grasshopper for potential herbivores. Ants (*Tapinoma sessile*, *Formica* sp.) were crawling on the rock and plants; however, we did not observe ants visiting axillary swellings, as would be expected for an extrafloral nectary producing sugary rewards. In addition, we did not observe any ant visitation to axillary swellings in the greenhouse-grown progeny placed next to peony plants.

### Observations from Growth Chamber Plant Culture

Since only one paper, the original species description, exists for *E. angulosa* (Nesom and Berger 2020), we include several observations from our experience growing this plant over the course of the study. Even with second-generation inbred seed of accession 1 and after stratifying for two weeks, germination was slow (2-3 weeks for the first germinants) and asynchronous, with new germinants continuing to emerge for several months and the robust morph germinating more slowly and at lower rates than the thin-stemmed morph. Floral longevity was extremely short, with corollas often falling off the plant within the same day of anthesis. Flowers readily self-pollinated without insect or hand-pollination, producing up to 50 seeds per fruit. In the growth chamber, plants readily produced stolons that rooted in the soil and hung over the side of the pot. Axillary swellings were often less conspicuous than those we observed in the field.

### Test for Genetic and Environmental Variation

In the test for effects of environmental and genetic variation on axillary swelling production, significant variance was attributable to family, but not to treatment (Family ICC: 0.732, Treatment ICC: 0.007). This result indicates that genetic variation for axillary swelling exists at the type locality.

Although treatment with JA, simulated herbivory, and soil amendments have been shown to induce extrafloral nectary size, number, and nectar secretion (Heil et al. 2001; Mondor and Addicott 2003; Pulice and Packer 2008), none of these treatments led to significantly increased axillary swelling. Instead, axillary swellings in our experiment were predicted by morph, with the thin-stemmed morph producing 83.7% (95% Confidence Interval (CI): [28.3%, 77.3%]) fewer swellings than the robust morph (P < 0.0001), and accession 1 producing 59.6% (95% CI: [77.9%, 87.9%]) fewer than the robust morph (P = 0.002, Fig. 2). We also observed no fluid secretion from any axillary swelling in this experiment, nor did glucose test strips detect any sugar production when we added drops of reverse osmosis water in an attempt to rehydrate any dried nectar.

**FIG. 2.**
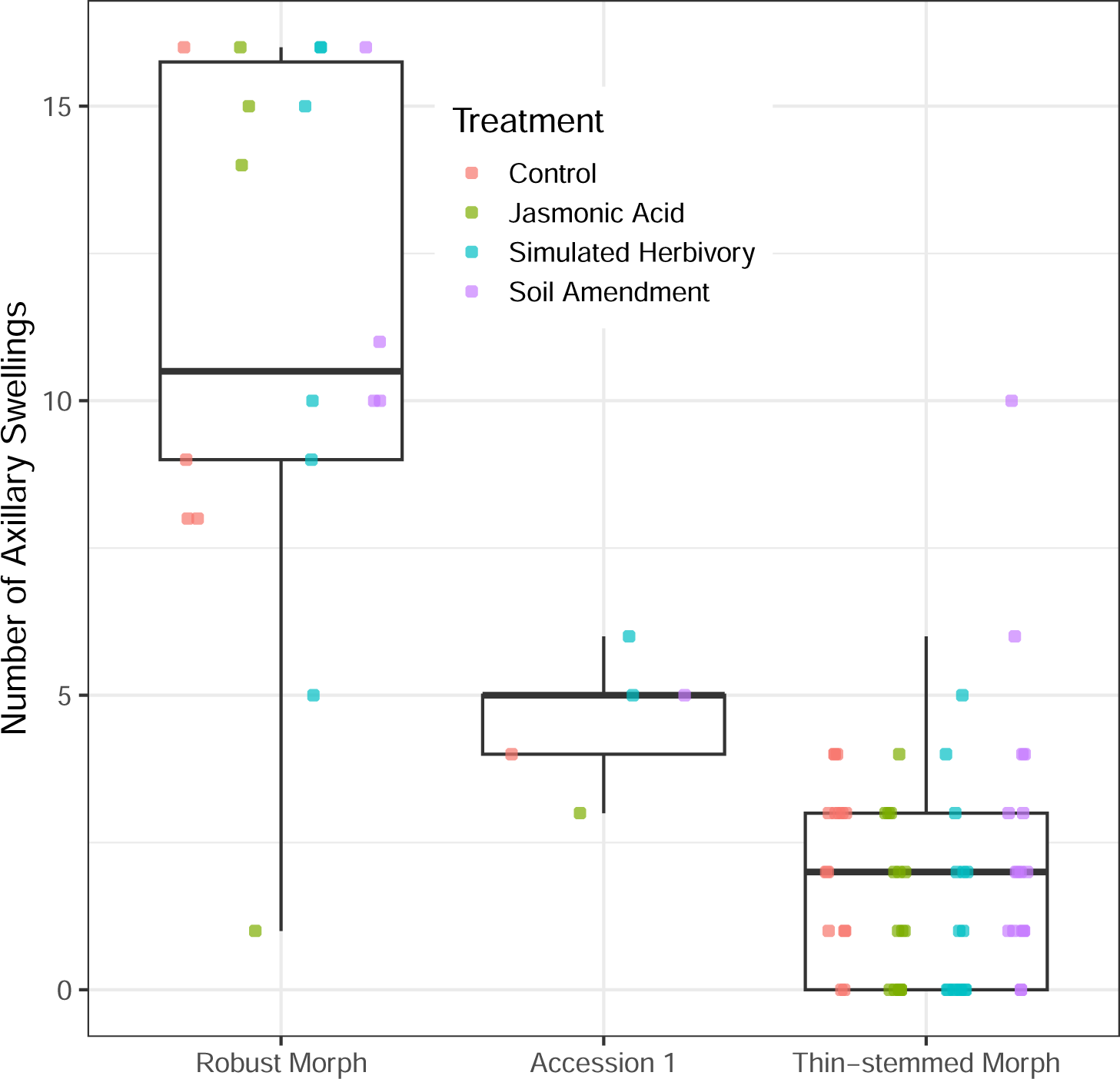
Observed variation in pedicel axillary swelling number across diverse germplasm collected from type locality. The robust morph had more axillary swellings than the thin-stemmed morph, with accession 1 intermediate between the two.

In addition to genetic variation for axillary swellings, we also found two distinct morphs with dramatic differences in their branch number, stem thickness, and leaf length (Fig. 3, Fig. 4). We observed (but did not measure) that the robust morph germinated slowly and had more glandular trichomes than the thin-stemmed morph, which germinates quickly and is glabrous. The robust morph log leaf length was 2.15 standard deviations (95% CI: [1.74, 2.55]) larger than that of the thin-stemmed morph (Fig. 4A, P < 0.0001). In the robust morph, log stem thickness was 2.44 standard deviations (95% CI: [2.18, 2.70]) greater than those of the thin-stemmed morph (Fig. 4B, P < 0.0001). In addition, thin-stemmed morphs had 2.49 times (95% CI: [1.95, 3.18]) more stems on average than the robust morph (Fig. 4C, P < 0.0001). However, there were 11.0% (95% CI: [7.92%, 14.0%]) fewer axillary swellings for every additional branch (P < 0.001), indicating that genetic variation for branch number does not drive axillary swelling variation. Moreover, the robust morphs were younger at phenotyping due to their delayed germination (see Observations from Growth Chamber Plant Culture), yet produced more axillary swellings than the thin-stemmed morph. Thus, the difference in age is unlikely to drive differences in axillary swelling number because axillary swellings were only seen on mature plants.

**FIG. 3.**
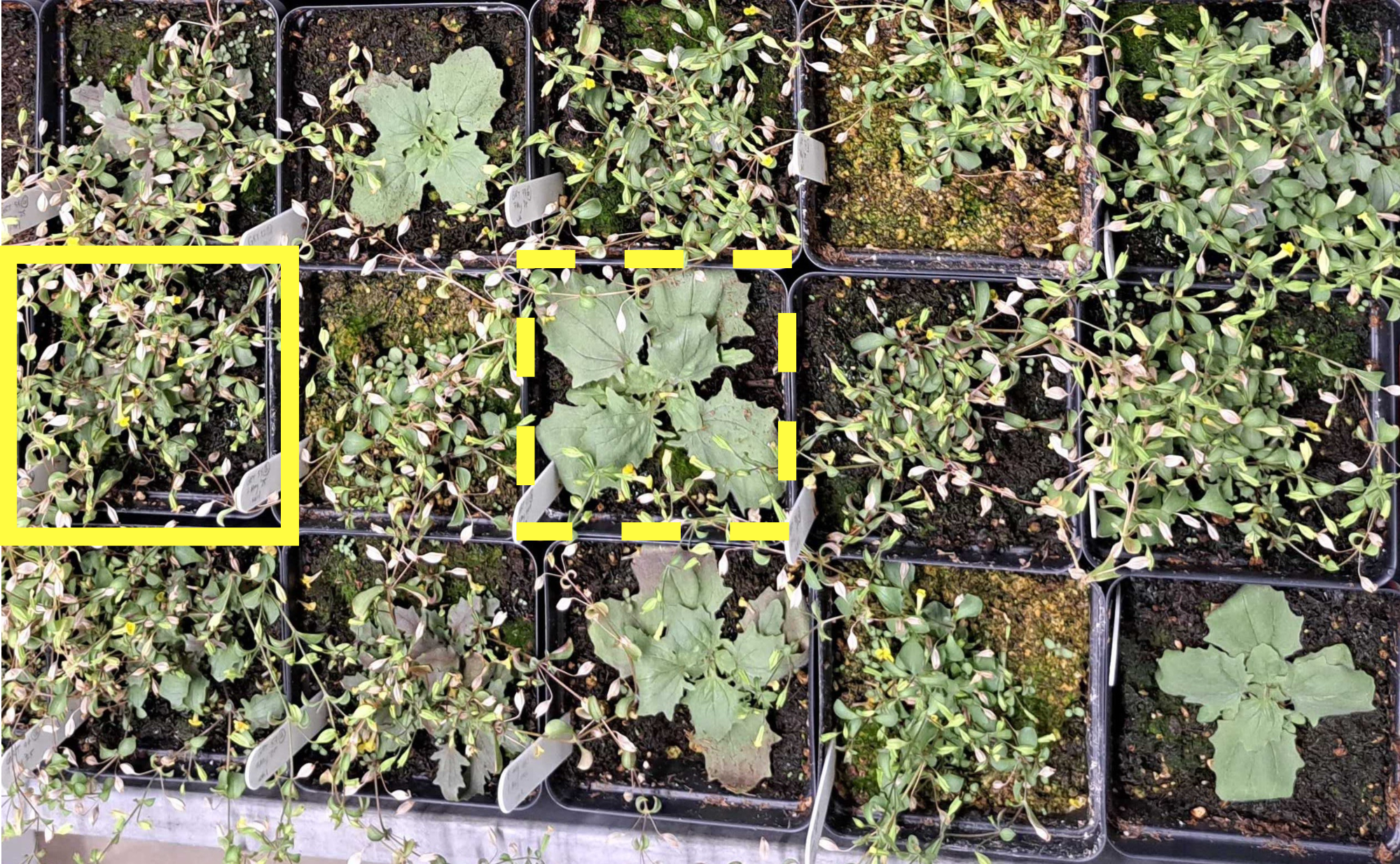
Thin-stemmed morph with many flowering branches and small, glabrous leaves (example in solid yellow box), and robust morph with few large leaves, many glandular trichomes, and less branching (example in dashed yellow box).

**FIG. 4.**
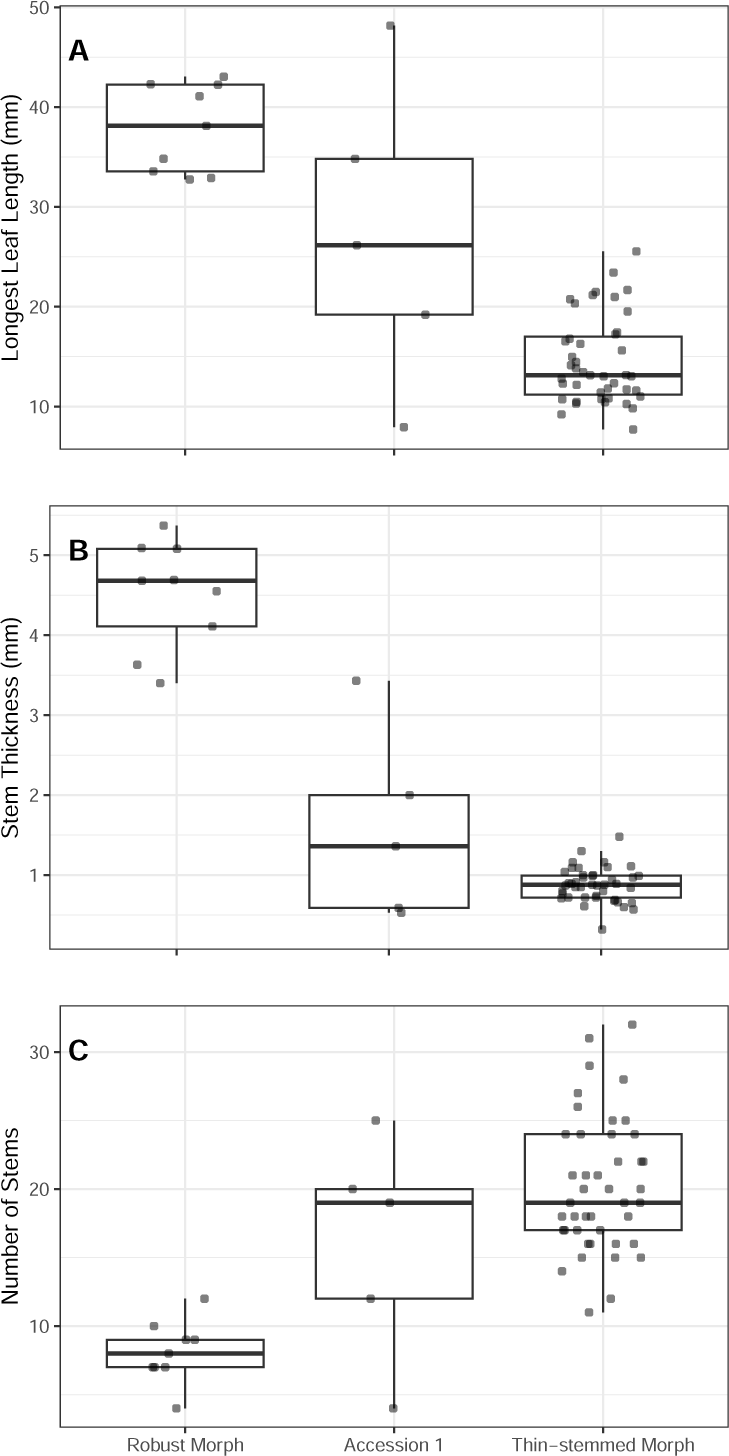
Morphological differences between the two morphs. (A) Robust morphs had larger leaves than thin-stemmed morphs; accession 1 took a wide range of leaf sizes falling intermediate to the two morphs. (B) The robust morph and thin-stemmed morphs had markedly different stem thicknesses, with Accession 1 again taking an intermediate phenotype. (C) The robust morph had many fewer stems than the thin-stemmed morph.

### Laboratory Observations of Crickets

We did not observe widespread targeted herbivory of axillary swellings by crickets in our feeding assay, in contrast to past reports (Gish et al. 2015) of targeted herbivory of fava bean EFNs. Out of 19 replicated herbivory assays, only a single cricket targeted the axillary swelling over leaf, petiole, or stem tissue.

### Microscopy

Scanning electron micrographs revealed that stomata are present on petioles (Fig. 5C) and on both sides of leaves, but scarce to absent on the axillary swellings themselves (Fig. 5A, 5B). Although we observed dense glandular trichomes on the junction between the swelling and the pedicel (Fig. 4B), these were not situated on the swelling itself, and glucose test strips did not detect glucose from these secretions. Moreover, various congeners produce glandular trichomes (Holeski et al. 2010; Bustamante Eguiguren et al. 2020) with no evidence for nectar secretion or a function in defense-mutualism. Under light microscopy, we observed that the axillary swellings are mostly composed of large-celled cortex tissue, with minimal incursion of vascular tissue into the swelling (Fig. 5D, 5E). In summary, microscopic analysis revealed no evidence of nectaries as modified stomata, some production of glandular trichomes near the swelling, and minimal vascularization of the swelling.

**FIG. 5.**
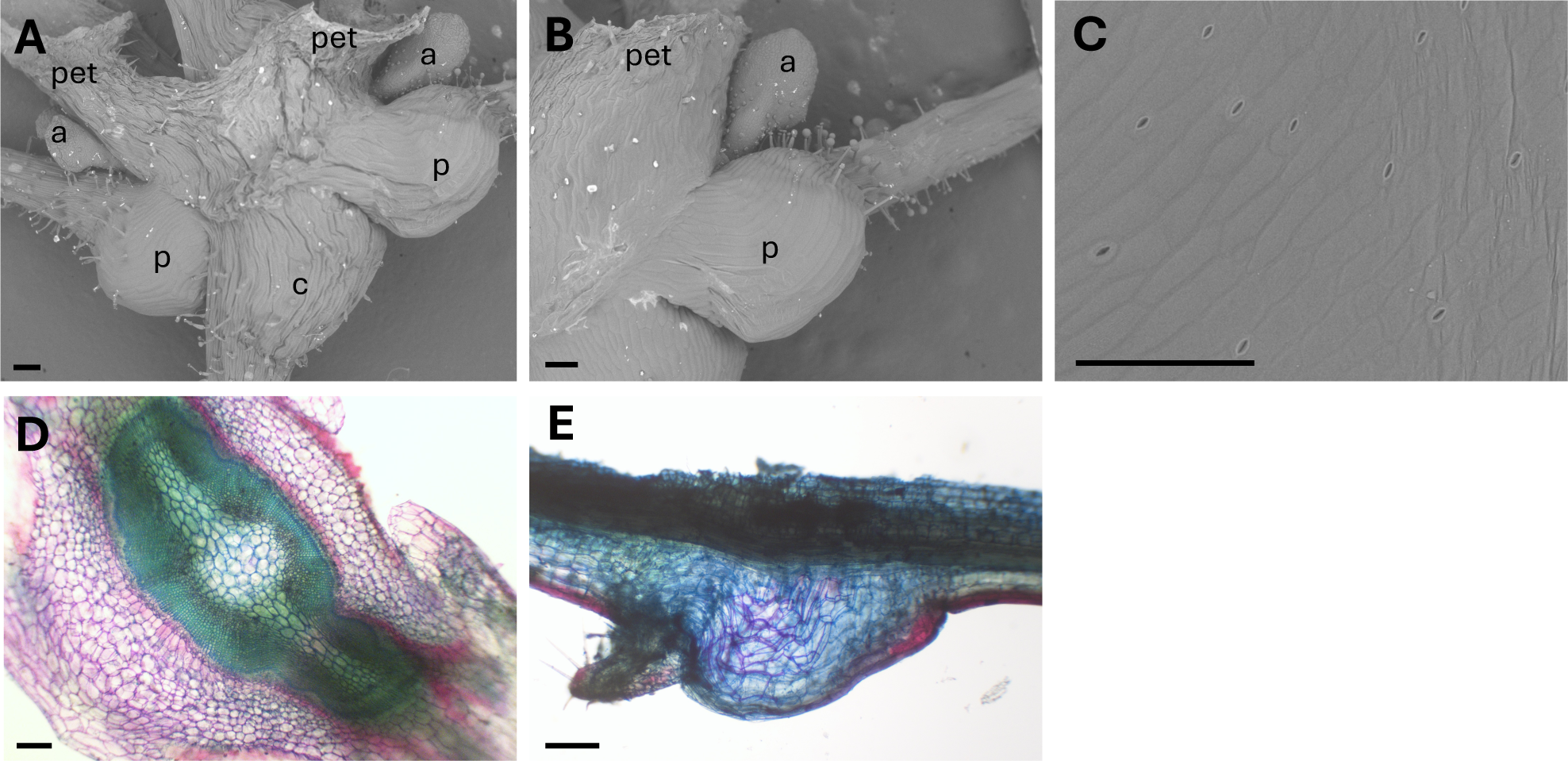
Morphology of axillary swelling showing (A) entire axillary swelling from SEM, (B) swelling at base of pedicel from SEM, (C) petiole with stomata visible from SEM, (D) cross section of axillary swelling through central swelling, stained with Toluidine Blue O under light microscope, (E) longitudinal section of axillary swelling through central swelling, stained with Toluidine Blue O under light microscope. Top of image is broken (through vascular tissue and pith) due to incomplete section. Abbreviations: a = axillary bud, c = central swelling, p = pedicel swelling, pet = petiole. All scale bars indicate 200 µm.

## Discussion

Here, we holistically investigate axillary swellings reported in *Erythranthe angulosa* for sugar-secreting function using a combination of field observations, laboratory experiments, insect bioassays, and microscopy. We found no evidence that this structure secreted sugar or functioned as an extrafloral nectary. In the course of our investigation, we uncovered two distinct morphs at the type locality that vary in shoot architecture traits and the number of axillary swellings. While we found no evidence for EFN production in *E. angulosa* in our study, we suggest these striking axillary swellings may serve a storage function, or indeed may not be adaptive.

Across anatomical and biochemical analysis, field observations, and insect bioassays we found no evidence for extrafloral nectar production in *E. angulosa*. In the field, axillary swellings were the most developed and evident, but we saw no nectar production on swellings even after bagging, nor did glucose test strips detect any sugars on the plant surface, even when water was added to rehydrate any sugar from dried nectars. We did not observe ants specifically or repeatedly visit axillary swellings at either the type locality or in our Michigan field study, suggesting these structures do not recruit ants or other predatory arthropods, and thus do not function as a defensive EFN. While some EFNs have no visible anatomical features to differentiate them from other leaf or stem tissue (Marazzi et al. 2013a), SEM and light microscopy did not reveal any characteristic nectar-exporting structures such as specialized glands or higher concentrations of stomata on the axillary swelling; indeed these swellings had lower densities of stomata than other parts of the plant. Glandular trichomes were present on *E. angulosa* stems and can function as EFNs by secreting sugar, but glandular trichomes are common in this clade and we detected no glucose in trichome secretions. Although we did observe great variation in axillary swelling production, we did not detect an induction of these structures or nectar secretion in response to jasmonic acid, simulated herbivory, or silica amendment, as would be expected for a plant defense trait and as has been observed in other EFN-bearing species (Heil et al. 2001; Pulice and Packer 2008; Gowton et al. 2025). We do not uncover evidence that these structures are EFNs in our study.

EFN physiology is complex, and it remains possible that these axillary swellings may function as nectaries. Some EFNs secrete nectar only at certain times of year or under very specific environmental conditions (Calixto et al. 2021). Nectar composition is frequently but not always high in glucose (Shenoy et al. 2012), and nectar that is composed predominantly of other carbohydrates (e.g. fructose) would produce a negative result on our glucose test strips.

However, none of our tests produced a visible drop of liquid, a hallmark of EFNs regardless of sugar composition. While it is still possible that this species produces extrafloral nectar only under conditions we did not test in our study, in our exhaustive search in the field and two growth chamber conditions, and with crickets, ants, glucose test strips, and microscopy, we found no evidence for extrafloral nectaries in *E. angulosa*.

Instead of serving as extrafloral nectaries, the axillary swellings may perform a different function. Our light microscopy analysis revealed that the axillary swelling is primarily cortex (parenchyma) tissue, with the vascular tissues extending minimally into the swelling (Figs. 5D, 5E). We propose that this expanded region of parenchyma may harbor starch reserves or store other metabolites, which could be tested with histological or metabolic analysis. Organs specialized for storage are usually located belowground, as in rhizomes of *E. corallina* growing nearby (Coughlan et al. 2021). However, large belowground reserves would not be possible in the granite rock outcrop habitat of *E. angulosa*, and rare cases of aboveground storage organs exist in nature, such as the air potato *Dioscorea bulbifera* (Celestine and David 2015). Alternatively, the axillary swelling and the many layers of xylem cells we observed may offer mechanical support or facilitate bending at the nodes, as the pulvini of the sensitive plant *Mimosa pudica* do for leaves (Sleboda 2023). Such a function may allow the plant to grow appressed to an uneven surface or respond to gravitational cues experienced by the hanging branches that produce abundant axillary swellings in *E. angulosa*. Stem swellings are sometimes associated with shoot-borne roots (e.g. Lenssen et al. 2000); however, the *E. guttata* species complex commonly produces shoot-borne roots without such swellings (Lowry and Willis 2010), so we consider this function to be unlikely. Finally, many developmental abnormalities of unknown function exist in plants and do not necessarily offer a selective advantage. In this small population, such a trait may rise to high frequency due to genetic drift rather than selection.

One unexpected finding of our study was striking variation among seeds germinated from soil collected from underneath the type individual. Using maternal families sampled from a fruit on a live *E. angulosa* plant and from the seed bank, we uncovered genetic variation affecting the number of branches and axillary swellings. Taken together with qualitative observations during our experiment, we have identified two contrasting growth forms: a thick-stemmed, less branched, slow-germinating form with many trichomes and axillary swellings, and a thin-stemmed, highly branched, faster-germinating form that is glabrous and has fewer axillary swellings. This dramatic variation was not observable in the field at the time of observation, and speaks to the genetic and morphological diversity that seed banks can harbor.

Our findings of two distinct morphs within only 100 mL of soil offer avenues for future exploration of life-history variation on small spatial scales. We observed huge morph divergence in stem thickness, which is widely used as a proxy for life history in monkeyflower, with perennials having thicker stems than annuals (Lowry and Willis 2010; Friedman et al. 2015; Zell et al. 2024). *E. angulosa* has not been observed year-round in the field to confirm its life history; however, the existence of genetic variation affecting stem thickness suggests that some genotypes produce a few stems that are energetically expensive to build and maintain, while others produce a large number of cheaper stems. The type locality is within 1000 m of *E. corallina*, a rhizomatous perennial monkeyflower, and reproductive isolation between *E. angulosa* and other members of the *Erythranthe guttata* species complex has not been tested. One possibility is that hybridization between *E. corallina* and *E. angulosa* could have produced the range of phenotypes observed at the *E. angulosa* type locality, with Horseshoe Meadows Road comprising a contact zone between annual and perennial members of the species complex.

## Conclusion

We found no evidence across ecological, anatomical, and biochemical investigations that axillary swellings on *E. angulosa* function as extrafloral nectaries. However, we uncovered remarkable genetic variation within the type locality for the number of these axillary swellings of unknown function, as well as for traits related to life history and shoot architecture. Further research into the genetic architecture of axillary swelling production and other shoot architecture traits, the functional morphology of the axillary swelling, and the population genetic structure of *E. angulosa* and nearby congeners will offer exciting insights on life history evolution and the origin of morphological diversity.

## Data Accessibility

All data and code associated with this study are available in a Zenodo archive (https://zenodo.org/records/23404519, doi: 10.5281/zenodo.23404518). Representative individuals of the robust morph, thin-stemmed morph, and accession 1 (grown in 16 h days in MSU growth chambers) were deposited in the herbarium at Michigan State University (catalog numbers MSC0299305, MSC0299306, and MSC0299307).

## Author Contributions

SME and MLP made equal contributions to the paper. SME and MLP designed the study, secured funding, conducted the fieldwork and wrote the original draft. SME performed the induction experiment and analysis and led the insect bioassays. MLP led the microscopy analysis. PAS assisted with germplasm curation, created herbarium vouchers, and made initial observations of the two morphs. All authors read and approved the final manuscript.

## Acknowledgments

We thank the Plant Resilience Institute Seed Grant Program at Michigan State University for funding. MLP and SME were supported by the NSF Graduate Research Fellowship Program (DGE-184-8739 and DGE-223-578). We thank Rosemary Glos for assistance with SEM imaging, Kathy Toll for consultation on methyl jasmonate, Nate Sanders for ant identification, and the Lowry lab and two anonymous reviewers for helpful comments. This is Kellogg Biological Station contribution no. 2447.

